# Open-Source DNA-Encoded Library informatics Package for Design, Decoding, and Analysis: DELi

**DOI:** 10.1101/2025.02.25.640184

**Authors:** James Wellnitz, Brandon Novy, Travis Maxfield, Shu-Hang Lin, Ivanna Zhilinskaya, Matthew Axtman, Tina Leisner, Eric Merten, Jacqueline L. Norris-Drouin, Brian P. Hardy, Kenneth H. Pearce, Konstantin I. Popov

## Abstract

DNA-encoded library (DEL) technology has become a powerful tool in modern drug discovery. Fully harnessing its potential requires the use of extensive computational methodologies, which are often available only through proprietary software. This restricts accessibility for small teams lacking robust informatics support, hindering the growth of the technology. Here, we present DELi, an open-source DEL informatics platform designed for library design, NGS decoding and calling, and enrichment analysis. DELi supports a simple and easy to understand configuration setup to present a straightforward user interface. To showcase its capabilities, we used DELi to design an in-house custom, benzimidazole-based DEL (UNC DEL006), and performed proof-of-concept selection experiments against Bromodomain-containing Protein 4 (BRD4). The DELi decoding and analysis modules identified top-performing compounds, leading to the off-DNA synthesis of UNC11951, which was confirmed as a nanomolar BRD4 binder via isothermal titration calorimetry (ITC) and differential scanning fluorimetry (DSF). These results demonstrate DELi as an effective tool for DEL design and analysis. Furthermore, its open-source nature will promote ongoing development and contributions from the DEL community to expand its applications and capabilities, making DEL technology more widely accessible.

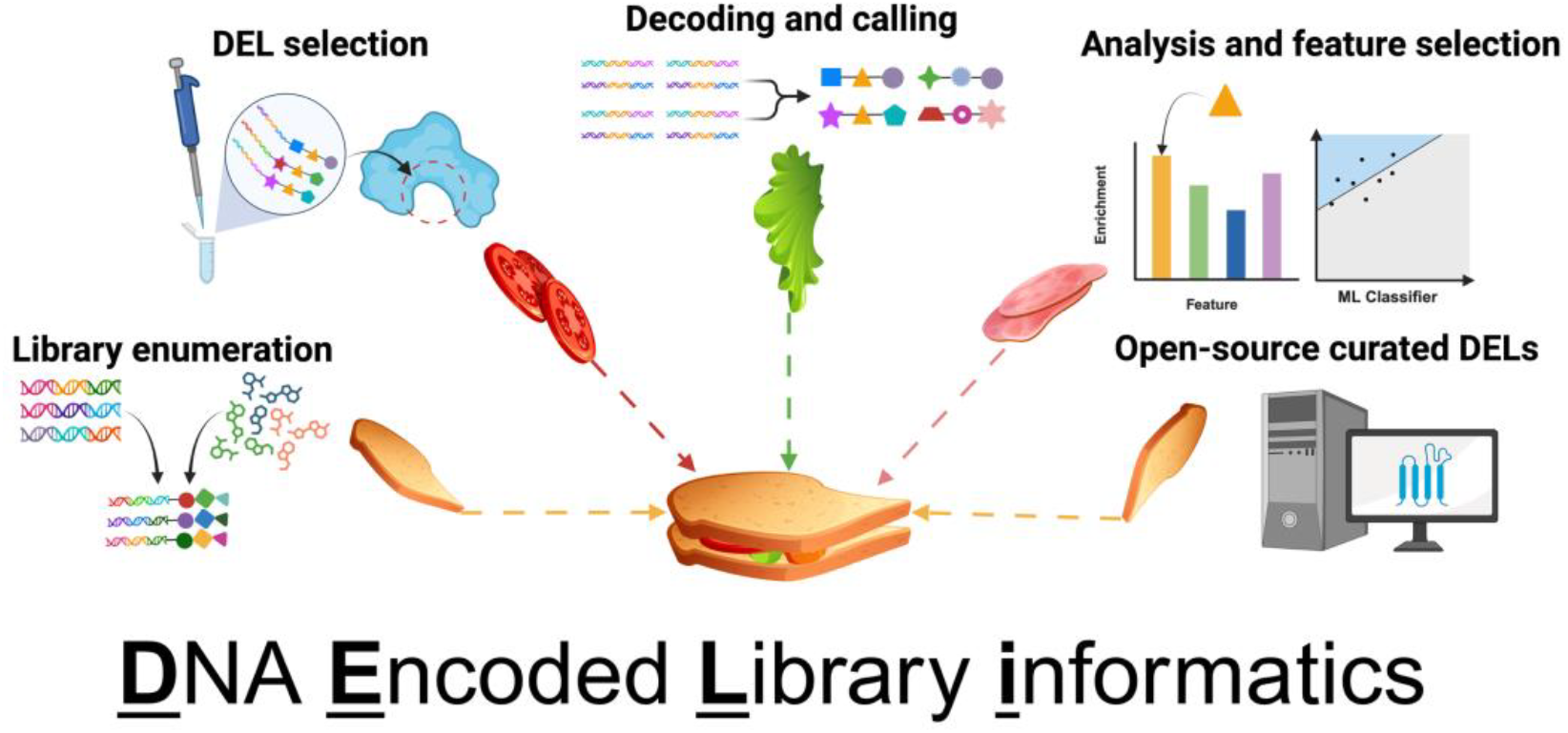

## Introduction

Drug discovery is a complex and expensive process, often costing over $1 billion USD and taking more than a decade to bring a therapeutic to market^1^. Technologies that improve efficiency while reducing costs are therefore essential for accelerating therapeutic development. Traditional high-throughput screening (HTS) has long been used to accelerate hit discovery, but its one-compound-per-well format is highly resource-intensive. To address this limitation, technologies such as phage and mRNA display were developed to enable the screening of millions to billions of compounds in a single tube^2^. While highly efficient, these methods are limited to peptides composed of natural or select unnatural amino acids^3^, constraining the chemical diversity of their libraries. DNA-Encoded Libraries (DELs) take advantage of the rapid screening capabilities pioneered by phage and mRNA display but overcome their chemical limitations by leveraging recent advances in combinatorial chemistry. As a result, DELs enable the rapid screening of billion-member libraries composed of structurally diverse, drug-like small molecules, significantly expanding the chemical space accessible to display technologies^4,5^.

Since their introduction in the 1990s^6^, DNA-Encoded Library (DEL) technologies have advanced rapidly, enabling the discovery of potent and selective small-molecule binders against a broad range of targets, including kinases^7^, GPCRs^8^, and histone-modifying proteins^9,10^. Commercialization by companies such as WuXi, HitGen, and Charles River Laboratories has made the screening of DEL libraries containing billions to trillions of compounds broadly accessible^11^. Moreover, to avoid the time- and resource-intensive process of resynthesizing DEL hits without DNA tags for confirmatory screening and prioritization, these companies increasingly use DEL screening data to train machine learning (ML) models. These models are then used to nominate purchasable hits by virtually screening commercial small-molecule libraries for compounds structurally similar to the original top DEL hits. Despite advances in screening, data processing and analysis continue to be major bottlenecks in the DEL workflow. These processes are often performed manually or using basic computational and visualization tools, which can introduces bias, slows down hit identification, and limits the ability to fully explore the vast chemical space encoded in DEL libraries.

DEL selection output consists of large volumes of DNA sequencing reads that require extensive processing to generate data that is both interpretable by humans and suitable for computational modeling. Converting raw sequencing reads into compound enrichment scores, a process known as decoding, involves statistical correction steps to account for noise introduced during sequencing, as well as variability arising from DEL synthesis and selection^12^. Despite the growing adoption of DELs, including open DELs^13^, the lack of open-source computational tools for data analysis remains a major barrier to accessibility. Smaller teams often lack the resources and expertise to implement published methods, many of which are not available as open-source software. In addition, the absence of internal informatics infrastructure further limits their ability to fully leverage the technology.

To address the lack of a powerful, flexible, and user-friendly open-source computational toolkit to support DEL technology, we developed the DNA Encoded Library informatics (DELi) software package. DELi offers a comprehensive and automated informatics pipeline, with modules that support DEL design, full library enumeration, selection decoding, and automated analysis of selection data. As an open-source academic initiative, DELi aims to make recent advances in DEL informatics widely accessible, providing a robust and extensible foundation to streamline and accelerate DEL-based research.

## Implementation

DELi is written in Python, a decision made to foster future collaborations and sustained support from the scientific community. The package encompasses all aspects of DEL informatics and is organized into separate modules to provide an intuitive user experience. These include `deli.decode`, `deli.enume`rate, `deli.analysis`, with each providing distinct capabilities for the varying steps of DEL informatics. Some functionalities are primarily accessed through a command line interface, while others are designed for use in custom scripts developed by users.

### DEL Configuration

A major barrier to creating an accessible, open-source DEL informatics package is the need for flexibility. DELs vary widely in design, synthesis routes, and DNA tag formats, requiring software that can accommodate diverse configurations while remaining easy to use. DELi addresses this challenge through a robust and highly configurable setup, supported by detailed documentation at both introductory and advanced levels. General users simply provide CSV or TSV files for building blocks and their mappings to DNA tags or SMILES, along with a concise JSON file that defines the library, including barcode structure and associated input files. These configuration files are typically under 50 lines, and DELi includes extensive syntax check that not only identifies errors but also suggests specific corrections, streamlining the setup process.

The modular architecture of DELi allows users to enable or disable specific modules depending on the task. For example, the `deli.analysis` module can be used independently for enrichment analysis and hit nomination, particularly when DEL screening and barcode decoding have been performed by a third party. This module remains functional even when chemical structures of DEL ligands are unavailable and only fingerprint information is provided. As with the automatic syntax checking feature, DELi can notify users early if a requested task requires missing information and can indicate what that information is, so it can be added when available.

### Error-Correcting Barcode Design

Error-correcting DNA barcodes are a well-established approach to reducing sequencing errors^14^, enabling recovery of up to 10 percent of total sequence reads^2^. The `design.barcode` module in DELi currently supports barcode design using a well-established Hamming encoding scheme^14^, which is intended to allow correction of single point mutations later during decoding. A custom quaternary Hamming encoder is used to generate barcode sets ranging from 7 to 16 base pairs, all guaranteed to have a minimum Hamming distance of three from every other member of the set. This enables SNP detection and correction using the corresponding Hamming decoder. An optional parity mode increases the minimum Hamming distance to four, allowing for detection, but not correction, of two SNPs, as well as correction of single SNPs. DELi does not yet support filtering nucleotide sequences based on GC content or repeated base pairs, but support for this feature is planned in the DELi roadmap (see below), along with tools for building block and DEL design.

### Library Enumeration

DELs are designed by combinatorially assembling sets of molecular building blocks through a common chemical reaction scheme. Computational enumeration of these building blocks into fully assembled molecules is an essential step for defining the chemical space represented by a library and can be a daunting task for large libraries. For example, 3,000 building blocks combined in two reactions can produce over a billion unique compounds. DELi provides integrated support for this enumeration process using user-defined design parameters specified in configuration files. If users supply the reaction scheme, DELi can perform the enumeration. Details on the required syntax are provided in the DELi documentation. DELi supports both batch enumeration of entire libraries and on-demand enumeration of individual compounds based on their unique identifiers. This functionality is also available through the command line interface. Future updates to DELi will include support for nonlinear and conditional synthesis schemes.

### Barcode Decoding

A core component of the DEL informatics pipeline is converting raw sequence reads collected after selection into compound counts for enrichment calculation. This involves matching the sequences to the DNA tags of the compounds that make up the screened library. DELi can decode heterogeneous collections of DELs. Regardless of the DNA barcode structure, library design, or the number of DELs used in each selection, DELi can perform decoding in a single run.

The decoding process begins with the user defining a DEL selection experiment in a human-readable YAML file, specifying the decoding settings and the DELs used. A single call to the `deli.decode` command-line interface then completes the decoding. On the back end, DELi uses a semi-global alignment algorithm implemented in cutadapt^15^ to map reads to their corresponding libraries. A second semi-global alignment, using a custom scoring matrix, is then applied to the full reference barcode to produce a robust mapping of each DNA read to the barcode schema. Both alignment steps support customizable error tolerance.

After alignment, DELi attempts to map the DNA sequence segments to the tags of the building blocks they encode. As noted above, DELi supports error-correcting tags, though their use is optional. When they are used, DELi provides two methods for error correction. If a valid Hamming code is used, DELi applies rapid error correction. To our knowledge, there are no open-source tools for generating tags with quaternary Hamming codes before DELi. As a result, many DEL barcodes were created through random generation rather than structured encoding. Because of this, DELs often used sets of barcodes that do not follow a single Hamming code scheme but instead lacked formal structure or combined multiple codes. To support these random sets, DELi uses a hash map lookup table for decoding. This method is equally fast but requires more memory, typically less than 100 megabytes.

After all building blocks are decoded, DELi will “count” the compounds. This counting process also supports unique molecule identifier (UMI) correction^16^, storing a separate count for just UMIs. Often both counts, raw and UMI corrected, are needed to calculate various enrichment metrics. After this, DELi writes the output counts for each decoded DEL compound to a CSV file, called a “cube” file in DELi.

The entire decoding process is tracked with robust logging in DELi, capturing detailed information about failure rates, including the specific reasons why a read could not be decoded (**Figure 1**). This information is summarized in a user-friendly, configurable HTML report. For well-calibrated DNA sequencers and validated DELs, DELi can successfully decode between 80-95% of reads, depending on the error correction scheme used by the library. It can process over one million reads per minute on a single CPU core. Parallelization is supported through Nextflow^17^, enabling efficient execution on high-performance computing clusters or distributed systems. Using DELi, we can decode 10 billion reads in under 15 minutes on an academic high-performance computing (HPC) system with approximately 800 cores. On Amazon Web Services (AWS) or Google Cloud Platform (GCP), the same task can be completed in under an hour at a cost of less than 10 USD.

**Figure 1:**
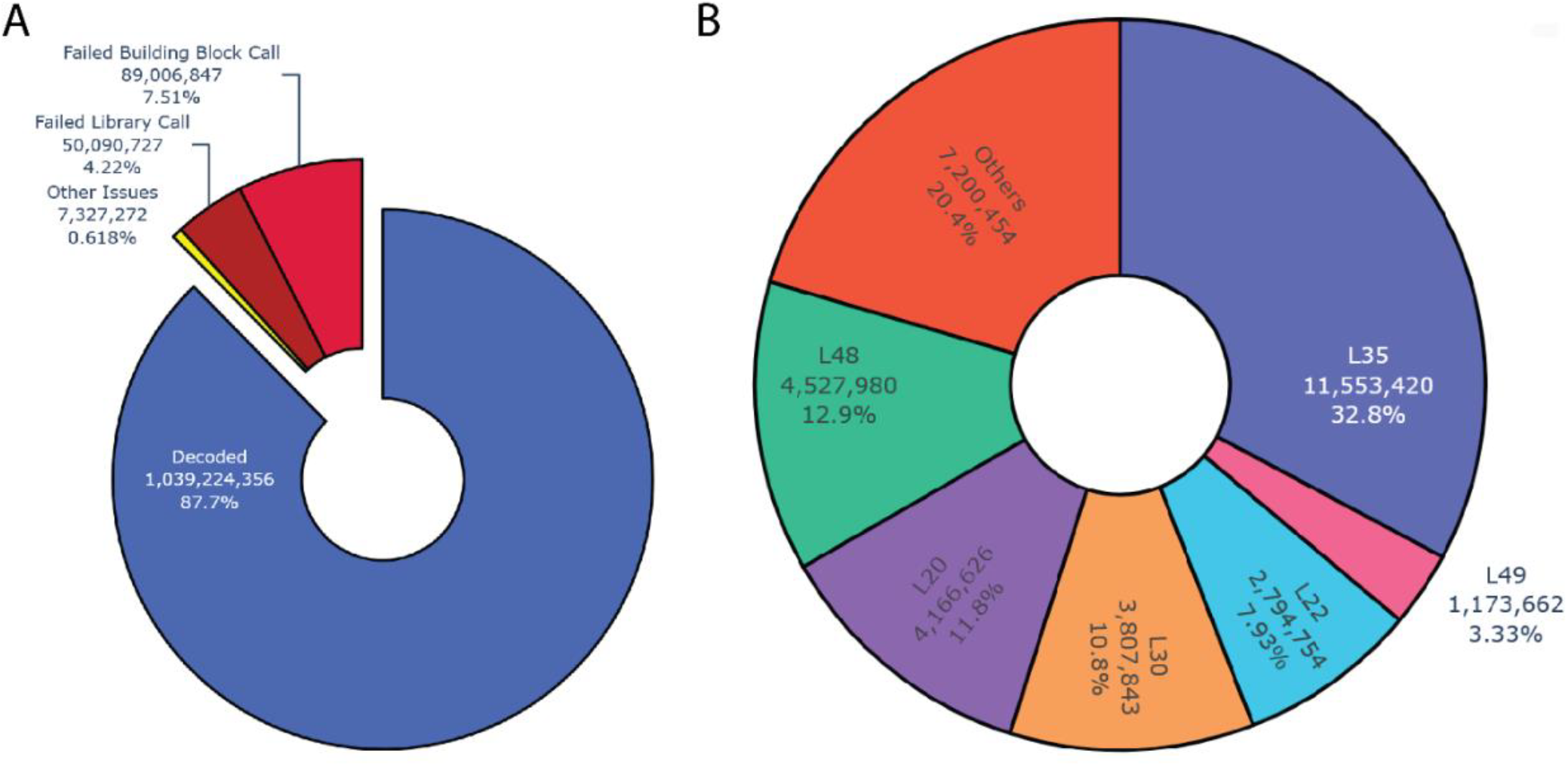
Sample graphs generated by the DELi decoding HTML report: A) Pie chart showing how many reads failed to be decoded and the relevant cause. B) Pie chart of which libraries were found in the selection and the percentages of UMI corrected counts attributed to that library (above a cutoff of 1%).

### DEL Analysis

Once decoding is complete, the resulting DEL selection data can be analyzed using the `deli.analysis` module to calculate enrichment scores for each compound. When multiple libraries are screened simultaneously, differences in library size and baseline noise can make it difficult to compare raw UMI-corrected counts directly. To address this, various enrichment metrics have been proposed in the literature^18^. DELi implements a suite of these methods, allowing users to choose the most appropriate metric for their specific experiment (Table 1). DELi also supports di/monosynthon analysis, a commonly used approach for identifying strong and consistent trends in DEL binding data. Enrichment metrics can be applied either at the fully enumerated compound level or at the synthon level, depending on the method used. In addition, DELi includes machine learning-based tools for quality control, helping to detect signals, assess data reliability, and further reduce noise in selection results.

**Table 1.** DEL Statistical Methods Available in DELi.

| Method | References |
| --- | --- |
| NGS Sampling Depth | McCarthy et al. (2020)<sup>31</sup> |
| Normalized Sequence Count | Franzini et al. (2015)<sup>32</sup> |
| Maximum-Likelihood Enrichment Ratio | Hou et al. (2023)<sup>33</sup> |
| Normalized Z-Score | Faver et al. (2019)<sup>21</sup> |
| PolyO | Chen et al. (2022)<sup>23</sup> |
| DEL-Based Random Forest | McCloskey et al. (2020)<sup>19</sup> |
| DEL-Based Graph Convolutional Network and Graph Attention Network | Duvenaud et al. (2015)<sup>34</sup> |

In addition to enrichment analysis, DELi offers a range of optional graphical visualizations to support feature selection. These include tools for analyzing competition binding experiments and rendering compound structures for structure-based selections (**Figure 2A and 2B**). DELi also incorporates automated data balancing functions to improve the performance of baseline machine learning models when modeling is desired. These models can be used for downstream virtual screening guided by DEL data^19^. Baseline models also serve as a diagnostic tool to assess whether a selection contains a detectable and “learnable” signal, providing an additional layer of quality control. DELi supports both classification and regression approaches (**Figure 2C**), streamlining model development and enabling more accurate and robust predictions in drug discovery workflows that utilize machine learning.

**Figure 2.**
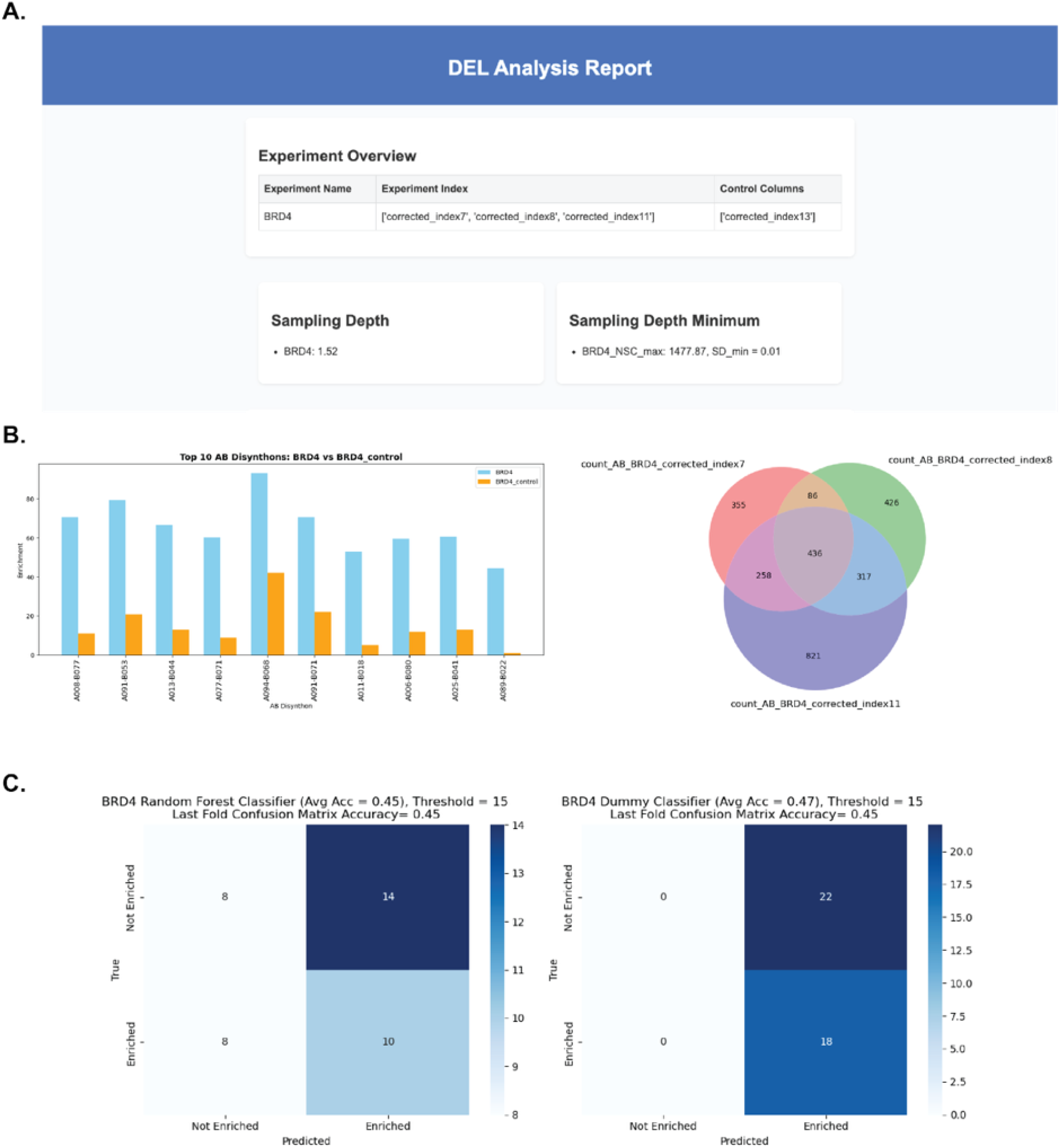
Automated DELi Analysis Report Accelerates Selection QC and ML Workflows. A) Header from DELi report detailing sampling depth and experimental conditions. B) Top AB-disynthon features from UNC DEL006, nominated by their enrichment over an NTC condition. The Venn diagram displays the AB-disynthon feature reproducibility across the three replicate selections, given a user-defined enrichment threshold. C) Automated DEL-ML RF classification model created by DELi’s data balancing functions contrasted with dummy classifier to display overall accuracy from 5-fold training regime.

### Parallelization

While DELi does not provide native parallelization within the package, most analysis metrics are inexpensive to compute after decoding. Although decoding itself can be computationally intensive, it is inherently well-suited to parallel processing. Different segments of the data can be decoded independently with minimal need for inter-process communication (“embarrassingly parallel”). As such, parallelization is left to the user to implement according to their system configuration.

One limitation of this approach is related to the UMI-corrected count calculation during decoding. Accurate correction requires coordination across parallel decoding jobs to track which UMIs have already been observed. To support this, DELi allows each decoder to save its UMI state as a JSON file upon completion. These files can then be merged using a simple script. To facilitate deployment, we provide a Nextflow workflow script that enables DELi decoding to be parallelized across distributed systems. This external workflow support allows DELi to be efficiently deployed on a wide range of infrastructure setups with only minor customization. The Nextflow workflow has been tested on HPC-SLURM clusters as well as in AWS and GCP cloud environments.

### Installation and Configuration

DELi is platform/operating system independent and made available for installation via Python pip. It can also be installed from source using Poetry for development purposes. Installation will automatically add DELi modules to the command line.

A portion of functionalities, like decoding and enumeration, require users to generate configuration files outlining the setup and contents of their DEL. Detailed documentation with examples is provided on how to generate these files.

### BRD4 Case Study with DEL6

To evaluate our DEL informatics pipeline, we used DELi to guide the design and synthesis of UNC DEL006, a benzimidazole based DNA encoded library (Figure S1). Leveraging a combination of commercially available and in-house building blocks, we employed DELi’s library enumeration module, `deli.enumerate`, to generate chemical structures and predict physicochemical properties of all possible DEL trisynthons. This enabled us to prioritize a set of A, B, and C position building blocks, 96 of each, for inclusion in the UNC DEL006 library. After selecting the chemical structures of the DEL ligands, we designed Hamming encoded barcodes for the three-cycle library. The final library was then synthesized using standard split-and-pool combinatorial methods. To further validate our computational workflow, we performed selection experiments against Bromodomain containing Protein 4 (BRD4), a well-characterized protein target implicated in cancer^20^. Experimental procedures for synthesis, selection, amplification, and sequencing are described in the Methods section.

We then performed barcode decoding using `deli.decode` and enrichment analysis using `deli.analysis` to identify top DEL hits by analyzing common features among the most enriched compounds at the trisynthon level, as well as through disynthon-based aggregation analysis. Both modules prepared and aggregated our replicate screening data into their respective module reports.

From the prioritized trisynthon compounds automatically reported in the DELi Analysis Report using a normalized z-score metric^21^, we selected candidates for off-DNA synthesis, follow-up characterization, and confirmatory screening (Figure 3A). We also utilized published chemical matter on our protein target to help in the prioritization stage^22^. Notably, one of the top-ranked hits nominated by DELi, UNC1195, was validated as a nanomolar binder of BRD4 using isothermal titration calorimetry (ITC) (Figure 3B). In contrast, a structurally-similar compound, UNC11954, which was not prioritized by DELi due to disynthon-level feature analysis, showed no detectable binding in the same assay. These results highlight DELi’s ability to distinguish active from inactive chemotypes by analyzing DEL selection data. Further validation using an orthogonal biophysical approach, differential scanning fluorimetry (DSF), confirmed the binding activity of UNC11951 (Figure 3B).

**Figure 3.**
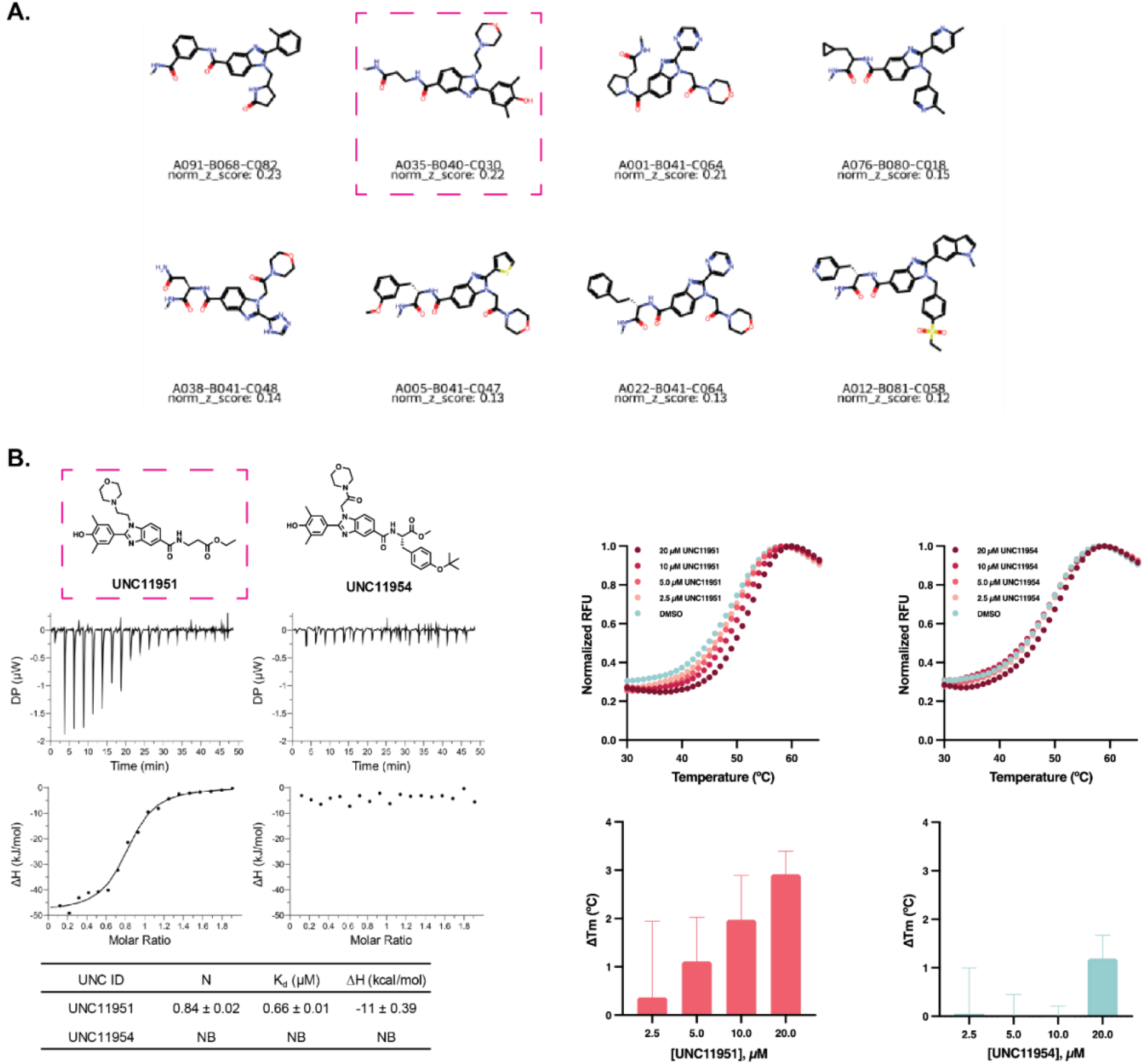
DELi Analysis Module Nominates nM Binder From DEL Selection. A) Top trisynthon compounds for SAR analysis utilizing a normalized z-score metric. B). ITC data for the top-nominated compound UNC11951, demonstrating nanomolar binding affinity, compared to the structurally similar UNC11954, which was not nominated by DELi’s automated report and showed no measurable binding affinity by ITC. C) Thermal shift assay results: melting curves (top) and calculated T_m_ values (bottom) for four-point dose-response experiments (20 to 2.5 μM) with UNC11951 and UNC11954. T_m_ values were determined by fitting the raw fluorescence data using the Boltzmann sigmoidal equation in GraphPad Prism. Error bars indicate the standard deviation of the calculated T_m_ values (n=3).

## Discussion

In developing DELi, we observed that a wide range of statistical methodologies have been reported in the literature that, in principle, could support robust analysis of DEL data^18^. However, many of these approaches were originally developed for other domains or lack implementations that are readily adaptable to DEL data. Considering this, the primary goal of DELi is not to introduce new analytical algorithms or computational frameworks. Rather, DELi is designed to serve as an accessible, flexible, and well-documented platform that lowers the barrier to entry for researchers adopting DEL technology and supports the application of existing methods within a unified informatics pipeline. In this vein, we have also supplied sample data and open-sourced our DEL6 library to aid teams in establishing their DEL informatic pipelines.

A central objective of DELi is to help establish standardized best practices for processing and interpreting DEL data. DEL experiments are inherently noisy, and computational analysis plays a critical role in distinguishing meaningful signals from background artifacts. The quality of data analysis can profoundly influence the outcome of a DEL campaign, yet there is limited guidance in the literature regarding recommended practices, let alone reproducible and well-documented implementations. A persistent challenge in the field is the absence of standardized terminology, which can unintentionally hinder communication and reproducibility. For instance, one group may use “disynthon”^19^ while another group refers to the same part of a compound as a “feature”^23^, potentially hindering cross-study comparisons and tool development. DELi was designed to serve as a foundational framework for DEL informatics, providing a structured, reproducible, and extensible platform to support consistent data processing and facilitate the sharing of both DEL screening results and new informatics methodologies. As the field evolves, DELi is positioned to integrate emerging analytical approaches and contribute to the maturation and standardization of DEL informatics.

### End-to-End Open-Source DEL

Despite the growing interest in open science, DEL technology remains largely inaccessible due to the proprietary nature of many existing platforms. Most large-scale DEL providers are commercial entities that impose substantial costs for access, and synthetic schemes for library construction are often not publicly disclosed. However, this landscape is beginning to shift, with some vendors making libraries more openly available and new collaborative initiatives emerging. For a truly open-source DEL ecosystem to take hold, it is essential that the computational infrastructure supporting DEL analysis is also open, transparent, and freely accessible. DELi was developed with this principle in mind, providing an open-source informatics platform that complements recent efforts to democratize DEL screening and supports the broader adoption of the technology within the academic research community.

The open-source nature of DELi is fundamental to its design and long-term vision, directly addressing a critical barrier in the field. Transparent and accessible computational tools that are engineered for generalizability are essential to advance DEL technology and make it more accessible to academic laboratories and smaller biotechnology companies. By unifying and democratizing established analysis methods within a cohesive platform, DELi enables more rigorous and reproducible DEL data interpretation. It supports key aspects of informed decision-making, including evaluation of experimental reproducibility, optimization of sampling depth, strategic incorporation of competitor binding experiments, and scaffold-based analyses to prioritize compounds for follow-up studies.

By open sourcing our complete analysis and DEL design pipeline, along with selected DEL libraries, we aim to address the current scarcity of openly available DEL software and datasets. This integrated approach establishes a foundation for community-driven development by promoting the use of standardized tools and facilitating more accessible data processing and library design. Rather than confining researchers to proprietary platforms, DELi offers a transparent and adaptable framework that gives users greater control over their DEL workflows. Through this commitment to open science, we seek to foster a collaborative ecosystem in which the sharing of data, methodologies, and tools accelerates innovation and broadens access to DEL technology.

### The Role of DELi in Advancing DEL-ML

A growing area of interest in the DEL field is the integration of machine learning (ML) methods into DEL data analysis, often referred to as the DEL-ML paradigm^24^. As ML techniques become increasingly embedded in drug discovery workflows, the synergy between DEL screening and predictive modeling holds significant potential. Recent studies have begun to explore both the use of DEL datasets for training ML models and the development of ML tools to enhance various stages of the DEL pipeline^19,24,25^. However, the predictive accuracy and reliability of any ML model are critically dependent on the quality, consistency, and proper annotation of the underlying training data. A key open question is whether DEL data, as it is currently generated and processed, is of sufficient quality to support robust ML applications. This challenge is exacerbated in the absence of accurate, transparent, and standardized tools for processing raw DEL data. DELi is designed to address this gap by providing a reproducible and well-documented framework for data curation and analysis. Without such tools, it becomes increasingly difficult to benchmark ML methods or assess the impact of data quality on model performance. We anticipate that DELi will play an essential role in enabling the development of reliable and effective DEL-ML tools for future drug discovery applications.

### Future Features

DELi has a well-defined development roadmap (see DELi Github Repo) aimed at expanding its capabilities to meet the evolving needs of the DEL community. A major focus of future work is the integration of an advanced suite of DEL design modules to streamline the generation of novel libraries, especially focused. One planned direction is the development of a structure-informed design module that leverages protein target information to guide building block nomination. This module will be designed to interface seamlessly with widely used docking and shape-based virtual screening platforms^26–28^, supporting target-driven DEL construction. Additional priorities include enhancing the flexibility of DEL configuration to support more complex library architectures, expanding built-in machine learning workflows for DEL-ML applications, refining the command-line interface, and improving containerization and default workflows for ease of deployment. As an open-source platform, DELi actively welcomes community input and contributions through a transparent and documented development process. By continuing to evolve in collaboration with the broader scientific community, DELi aims to serve as a robust, extensible foundation for the next generation of DEL informatics and drug discovery research.

## Conclusion

The field of DEL has rapidly expanded in recent years, with a surge in studies reporting novel DEL libraries, screening targets, and selection strategies^29^. Ready-to-purchase DELs have become increasingly available to academic labs and small biotech companies seeking to integrate this powerful technology into their drug discovery efforts^13,30^. However, many of these libraries require proprietary software licenses that limit flexibility and customization, leaving researchers constrained by closed systems. To address this, we introduce DELi, an open-source platform with fully accessible code and pipelines, available on GitHub for implementation and collaboration. Our goal is to provide researchers with a transparent and adaptable toolset, enabling greater control over their DEL workflows. We welcome feedback from the computational community and are committed to expanding DELi’s capabilities, including the expansion of deep learning models to explore novel, non-DEL-like chemical spaces for drug discovery.

The field of DELs has expanded rapidly in recent years, with a growing number of studies reporting novel library architectures, screening targets, and selection methodologies. Commercially available DELs are increasingly accessible to academic laboratories and small biotechnology companies, offering opportunities to incorporate this powerful technology into early-stage drug discovery efforts. However, many of these platforms rely on proprietary software, limiting flexibility, transparency, and reproducibility. To address this gap, we present DELi -- an open-source informatics platform with fully accessible code and modular pipelines, available on GitHub for community use and contribution. Our aim is to empower researchers with a customizable and extensible framework that enables greater control over DEL data processing and analysis. We actively welcome feedback from the broader computational chemistry community and are committed to expanding DELi’s capabilities, including the integration of deep learning models to explore chemically diverse, non-traditional DEL spaces in support of next-generation drug discovery.

## Experimental Methods

### Protein expression and purification

The bromo domain of BRD4 (residues 44-168 of NP_001366220) was expressed with an N-terminal His-tag in a modified pET28 expression vector. The BRD4 expression construct was transformed into Rosetta BL21(DE3)pLysS competent cells (Novagen, MilliporeSigma). Protein expression was induced by growing cells at 37°C with shaking until the OD_600_ reached ~0.6 at which time the temperature was lowered to 18°C and expression was induced by adding 0.5mM IPTG and continuing shaking overnight. Cells were harvested by centrifugation and pellets were stored at −80°C.

BRD4 protein was purified by resuspending thawed cell pellets in 30ml of lysis buffer (50mM sodium phosphate pH 7.2, 50mM NaCl, 30mM imidazole, 1X EDTA free protease inhibitor cocktail (Roche Diagnostics) per liter of culture. Cells were lysed on ice by sonication with a Branson Digital 450 Sonifier (Branson Ultrasonics) at 40% amplitude for 12 cycles with each cycle consisting of a 20 second pulse followed by a 40 second rest. The cell lysate was clarified by centrifugation and loaded onto a HisTrap FF column (Cytiva) that had been preequilibrated with 10 column volumes of binding buffer (50mM sodium phosphate pH 7.2, 500mM NaCl, 30mM imidazole) using an AKTA FPLC (Cytiva). The column was washed with 15 column volumes of binding buffer and protein was eluted in a linear gradient to 100% elution buffer (50mM sodium phosphate pH 7.2, 500mM NaCl, 500mM imidazole) over 20 column volumes. Peak fractions containing the desired protein were pooled and concentrated to 2ml in Amicon Ultra-15 concentrators 10,000 molecular weight cut-off (Merck Millipore). Concentrated protein was loaded onto a HiLoad 26/60 Superdex 75 prep grade column (Cytiva) that had been preequilibrated with 1.2 column volumes of sizing buffer (25mM Tris pH 7.5, 250mM NaCl, 0.5mM 1mM DTT, 5% glycerol) using an ATKA Purifier (Cytiva). Protein was eluted isocratically in sizing buffer over 1.3 column volumes at a flow rate of 2ml/min collecting 3ml fractions. Peak fractions were analyzed for purity by SDS-PAGE and those containing pure protein were pooled and concentrated using Amicon Ultra-15 concentrators 10,000 molecular weight cut-off (Merck Millipore).

### BRD4 DNA-encoded library selection

DEL library selections were performed using IMAC PhyTip® 200+ tip columns (5 µl resin volume, Biotage) and a semi-automated pipettor (E4 XLS, Rainin). Prior to His-BRD4 capture, tips were equilibrated with selection buffer [20 mM HEPES, pH 7.5, 100 mM NaCl, 0.01% Tween-20, 0.2 mg/ml BSA (Millipore/Sigma), 0.2 mg/ml sheared salmon sperm (SSS) DNA (Sigma)] at 250 µl per min flow rate (3 cycles). His-BRD4 was diluted to 1.2 µg/µl (60 µl total volume) in selection buffer and captured onto IMAC tips in triplicate by continuous pipetting for 30 min at 250 µl per min flow rate. The BRD4 IMAC tips were washed 3 times with 150 µl selection buffer and immediately transferred to 10 pmol UNCDEL006 diluted in 50 µl selection buffer. BRD4 IMAC tips were incubated 1 h at room temperature with continuous pipetting. To control for non-specific binding, one IMAC tip without BRD4 was processed in parallel with the BRD4 captured tips. The tips were then washed 2 times with 150 µl selection buffer and 1 time in selection buffer without SSS DNA. To elute bound library molecules, the tips were incubated 10 min with continuous pipetting in 60 µl selection buffer (without SSS DNA) heated to 80°C. A second round of selection was performed by incubation of freshly prepared IMAC BRD4 tips (and control no target tip) with the eluted library molecules (supplemented with 0.2 mg/ml SSS DNA) as described above. Library molecules were eluted in 50 µl Tris-HCl, pH 7.5 buffer supplemented with 0.001% Tween-20. Samples from both rounds of selection were amplified by qPCR using unique identifier index primer sequences followed by PCR cleanup (GeneJet, Sigma) and agarose gel analysis. Samples were prepared for Nanopore NGS according to manufacturer instructions for amplicon sequencing (SQK-LSK114 ligation sequencing kit, Oxford Nanopore Technologies).

Briefly, 80 fmol of each PCR product was pooled and subjected to NEBNext Ultra II end repair/dA-tailing (New England Biolabs) and Nanopore adapter ligation prior to loading onto a MinION Flow Cell (R10.4.1, Oxford Nanopore Technologies).

### Isothermal titration calorimetry

ITC experiments were performed at 25 °C using a MicroCal PEAQ-ITC calorimeter (Malvern Panalytical, UK). BRD4-BD1 protein (20 µM) and compound (200 µM) were prepared in buffer containing 25 mM Tris (pH 7.5), 150 mM NaCl, 2 mM β-mercaptoethanol, and 1% (v/v) DMSO. The titration protocol consisted of a single initial injection of 0.2 µL compound into the sample cell, followed by 18 injections of 2 µL each. Injections were spaced 150 s apart with an injection duration of 4 s. The reference power was set to 10 cal/s. The first injection was excluded from data analysis. Titration data were analyzed using the MicroCal PEAQ-ITC Analysis Software (Malvern Panalytical, UK) and fit to a one-site binding model.

### Differential scanning fluorometry

BRD4-BD1 protein (1 µM) was prepared in 50 mM Tris-HCl (pH 8.0), 200 mM NaCl, 1 mM DTT, and 1% (v/v) DMSO. Compounds were serially diluted 2-fold in the same buffer and pre-incubated with BRD4-BD1 at room temperature. After pre-incubation, SYPRO Orange Protein Stain (Invitrogen) was added to a final concentration of 5×. Fluorescence was monitored over a temperature gradient of 1 °C/min from 25 to 90 °C using an Analytik Jena qTower^3^ real-time PCR thermal cycler. Melting curves were analyzed in GraphPad Prism using a Boltzmann sigmoidal fit to determine the protein melting temperature (T_m_). Compounds were tested in triplicate and compared to a DMSO-only control. Thermal shifts (ΔT_m_) were calculated relative to the control.

## Acknowledgements

We thank the members of the Popov Lab and the CICBDD at UNC for helping to develop and giving feedback on DELi. We thank Valeriia Kaneva for valuable discussions during our python package preparation and testing. BN gratefully acknowledges support from the NIH Biophysics Training Grant (T32GM148376-01A1).

## Availability of data and materials

DELi code and instructions for installation can be found on our GitHub repo at https://github.com/Popov-Lab-UNC/DELi. Our open-source DEL6 library is made available as a CSV file. We also provide example selection data for testing the analysis module.

## FIGURES AND TABLES

**Figure S1.**
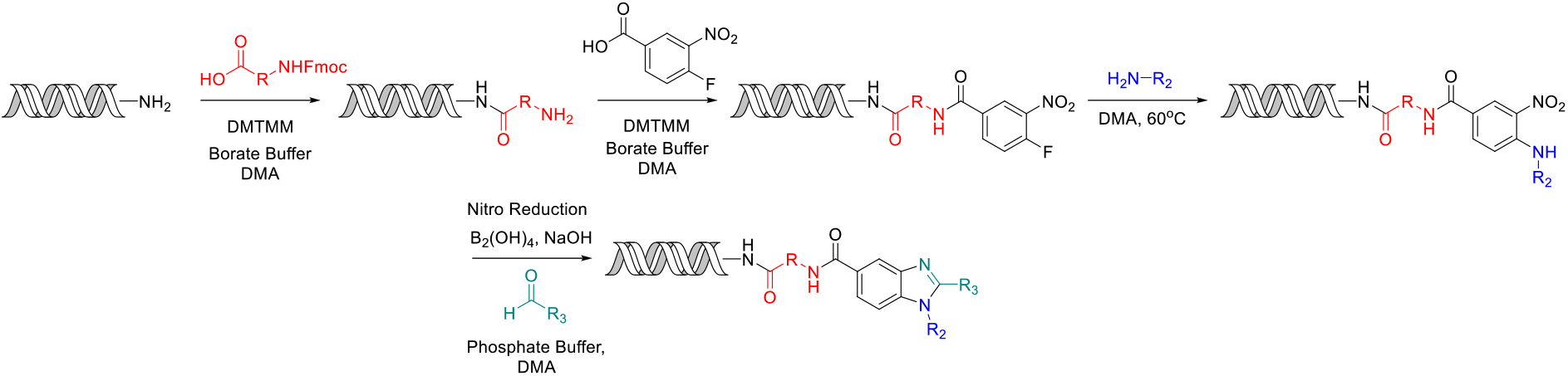
UNC DEL006 Synthesis.

## S1. DEL Synthesis Procedure

### General Procedure of Acylation of HP

To a 1mM (2000 uL) solution of HP in 250 mM Borate Buffer (pH9.5) was added 40 equivalents of acid (440 uL of 200mM solution, 49.2 mg into 504 uL of DMA) followed by 35 equivalents of DMT-MM (390 uL of 200mM solution, 29.4 mg into 531 uL of Borate Buffer). The reaction was allowed to proceed at room temperature with gentle mixing overnight. At the completion of the reaction, the DNA was crashed out by addition of 10% total volume of 5M NaCl (283 uL) followed by 3x total volume of cold EtOH (9339 uL). This was placed in a −80C freezer for approximately 30 minutes then centrifuged at 4C to give pellet.

### General Procedure for Nucleophilic Substitution

Resuspend the pellet to a 1mM solution with Borate Buffer and transfer 5 nmol of HP to 96 well plate to which 160 equivalents of amine were added (2.0 uL of 400mM solution). This mixture was heated to 60C for 16 hours. The DNA was then crashed out as previously described.

### General Procedure for Nitro Reduction

Resuspend the pellets to a 1mM solution in water and pool into a single reaction flask to which 500 equivalents of NaOH (43.2 uL of 5 M NaOH solution) was added followed by 1130 uL of EtOH and finally 150 equivalents of B_2_(OH)_4_ (648 uL of 100mM solution) was added and allowed to react for 2 hours. The DNA was then crashed out a previously described, twice.

### General Procedure for Benzimidazole formation

Resuspend the pellets to a 1mM solution in Phosphate Buffer (250 mM, Ph 5.5) was aliquoted back into a 96 well plate to which 60 equivalents of aldehyde (1.5uL of a 200mM solution in Acetonitrile) was added. This was allowed to react overnight. The DNA was then crashed out as previously described.

### Ligation of DNA Barcodes

To a 1145uM solution of AOP-HP (19 uL in water) was split into 96 wells (150mM final concentration). 50.4 uL of a 500 uM Building block tag was added to each well (180 uM final concentration) followed by 10x T4 Ligase Buffer (14.0 uL), Water (53.8 uL) and finally T4 Ligase (8000 units/well, 2.8 uL). This was allowed to react at room temperature overnight. Ligations were confirmed by gel analysis and the DNA was crashed out as described above.

## S2. Off-DNA Synthesis

### ethyl 3-(4-fluoro-3-nitrobenzamido)propanoate (A)

4-fluoro-3-nitrobenzoic acid (104.2 mg, 1 Eq, 562.9 μmol) was dissolved in DMF (2.815 mL) to which ethyl 3-aminopropanoate (131.9 mg, 2 Eq, 1.126 mmol) was added. 2-(3H-[1,2,3]triazolo[4,5-b]pyridin-3-yl)-1,1,3,3-tetramethylisouronium hexafluorophosphate(V) (428.1 mg, 2 Eq, 1.126 mmol) was then added to the stirring solution and finally N-ethyl-N-isopropylpropan-2-amine (291.0 mg, 392 μL, 4 Eq, 2.252 mmol) was added dropwise and allowed to react overnight. The reaction was quenched by addition of water, extracted 3x with EtOAc, organics were washed with brine 2x, and solvents were removed to give crude material. TLC and LCMS show good conversion to desired product, so it was purified via normal phase chromatography using EtOAc/Hexanes to give desired product (86% yield).

### ethyl 3-(4-((2-morpholinoethyl)amino)-3-nitrobenzamido)propanoate (B)

Ethyl 3-(4-fluoro-3-nitrobenzamido)propanoate (46.0 mg, 1 Eq, 162 μmol) was dissolved in DMF (809 μL) to which 2-morpholinoethan-1-amine (42.1 mg, 2 Eq, 324 μmol)was added slowly. This reaction was then heated to 60 °C and allowed to react overnight. The reaction was quenched by addition of water, extracted 3x with EtOAc, organics were washed with brine and solvents were removed to give crude material. TLC and LCMS show complete conversion of starting material to desired product, but the LCMS shows an additional more polar peak that could correspond to the amine. The reaction was purified via normal phase chromatography (DCM/MeOH) to give the desired product (73% yield).

### ethyl 3-(2-(4-hydroxy-3,5-dimethylphenyl)-1-(2-morpholinoethyl)-1H-benzo[d]imidazole-5-carboxamido)propanoate (UNC11951)

Ethyl 3-(3-amino-4-((2-morpholinoethyl)amino)benzamido)propanoate (43.0 mg, 1 Eq, 118 μmol) was dissolved in DMF (787 μL) to which 4-hydroxy-3,5-dimethylbenzaldehyde (35.4 mg, 2 Eq, 236 μmol) was heated to 60 °C overnight. The material was quenched by addition of water, extracted 3x with EtOAc, combined organics were washed with brine, dried with sodium sulfate and the solvents were removed to give crude material. It was purified via reverse phase chromatography to give desired product (68% yield).

## S3. Analysis Background

We employ a suite of analytical techniques that attempt to identify trends in the target-enriched synthons and fully enumerated compounds. One such method is the normalized sequence count (NSC)^32^, which is functionally analogous to RPKM/TPKM^35^ in the RNA-seq literature or sequencing depth-based normalization in ChIP-Seq/ATAC-Seq experiments. The NSC normalizes the reads for a given DEL member by the sampling depth for that experiment where c_i_ represents the observed count for a library member and SD is the sampling depth for a given target (Eq. 1). One benefit of the NSC is that it doesn’t require a separate control experiment or naïve sequencing run—thus effectively cutting the costs and accessibility to conduct DEL experiments.

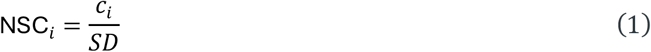

Using this formulation of NSC, we calculate the merged maximum-likelihood enrichment ratio as proposed by Hou et al^33^. with a smoothing factor to account for inherent variance in DEL sequencing counts. Here c_1_ and c_2_ represent counts for a given library member from selection and control experiments respectively, while n represents total sequencing counts for that selection.

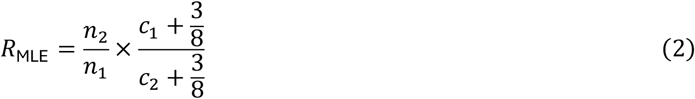

While users can provide DEL data without replicate samples, we opted for the merged calculation of the MLE ratio to increase confidence and raise the overall sequencing floor^33,36^. The normalized Z-score implemented by Faver et al^21^. models DEL selection data using a binomial distribution, which describes the probability of observing a given compound (or synthon/disynthon) x times across n independent trials with replacement. Here p_o_ represents the observed probability, p_i_ is the expected probability, and c_i_ are control counts for the given library member.

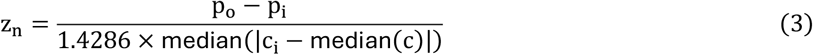

DELi also implements HitGen’s PolyO score^23^ for disynthon/monosynthon feature selection. This approach establishes a baseline score based on sequencing depth and size of a given DEL, then calculates the fold-change from the established baseline to determine if a feature is enriched.

